# Autoreactomes in Healthy Individuals Vary According to HLA Class II Genotype

**DOI:** 10.64898/2026.02.19.706904

**Authors:** Rayo Suseno, Juliano A. Boquett, Ravi Dandekar, Thomas Ituarte, Akshay Sharathchandra, Bonny Alvarenga, Cynthia Vierra-Green, Stephen Spellman, Martin Maiers, Joseph L. DeRisi, Michael R. Wilson, Jill A. Hollenbach

## Abstract

The link between *HLA* genotype and the presence of pathogenic autoantibodies has previously been established across different autoimmune disorders. However, the functional process linking specific antibodies to specific *HLA* remains unclear. Additionally, autoantibodies – usually associated with autoimmune disease – are also present in healthy individuals. To this end, we sought to determine the spectrum of self-antigen antibody specificity (the “autoreactome”) in healthy individuals, stratified by *HLA* genotypes. We utilized Phage ImmunoPrecipitation Sequencing (PhIP-Seq), a programmable phage display for interrogation of antibody specificity, which encompasses over 700,000 peptides tiled across the entire human proteome. Serum from 741 donors without diagnosed autoimmune disease were grouped by their five homozygous *HLA-DRB1* genotypes and analyzed for differences in autoreactivity profiles. We applied a custom filter to the PhIP-Seq normalized data and obtained a set of enriched peptides for each *HLA-DRB1* genotype. We found that the autoreactome in healthy individuals with different *HLA-DRB1* genotypes are generally distinct. Binary logistic regression successfully identified whether a sample belongs to a specific *HLA-DRB1* genotype (*HLA-DRB1*01, HLA-DRB1*03, HLA-DRB1*04, HLA-DRB1*07, and HLA-DRB1*15*) with the following accuracies: 96%, 92%, 90%, 94%, and 90%, and multinomial models predicted *HLA-DRB1* genotype with up to 90% accuracy. Finally, gene-level analysis suggests that individuals with specific *HLA* autoimmune risk alleles may harbor potentially pathogenic autoantibodies in the absence of, or prior to the establishment of, overt disease. Our analysis demonstrates that autoreactivity profiles in healthy people vary according to *HLA* class II genotype, and may provide insight into the pathological processes associated with development of autoimmune and other immune-mediated diseases.

## Introduction

Located on human chromosome 6, Human Leukocyte Antigen (*HLA*) genes encode cell surface receptors that present antigens to T cells, playing a pivotal role in the adaptive immune response^1^. *HLA* loci are the most polymorphic of the human genome, and this remarkable variation leads to distinctive peptide-binding repertoires, mediating key differences in antigen presentation and differentially eliciting immune responses^2^. Accordingly, *HLA* variation has been associated to a multitude of diseases, ranging from autoimmune, infection, neurological, to cancer phenotypes^3–8^.

Patterns of antibody response and the specificity of antibody epitope recognition in both infectious and autoimmune disease have driven the search for antigen bound by *HLA* and presented to T cells, in the interest of exploration of disease triggers, therapeutics, and vaccine development. B cells function as professional antigen presenting cells and are capable of accessing antigen for phagocytosis through the B cell receptor (BCR), a bound immunoglobulin (Ig). Evidence suggests that the BCR itself may be presented by *HLA* molecules and therefore be recognized as epitopes by T cells^9^. Data also suggest that there exists operative T-cell-mediated immunosurveillance with respect to follicular lymphoma cells, and that this control is mediated by the strength of binding of BCR-derived peptides to *HLA*^9^. Thus, the peripheral BCR repertoire may be shaped by *HLA* alleles in healthy individuals and controlled by T-cell mediated recognition of BCR peptides.

It has been shown that *HLA* class II modulates antibody response to both viral and bacterial antigens in vaccination^10–13^, as well as specificity for viral epitopes^14^. Furthermore, there is evidence for a link between *HLA* class II genotype and the presence of specific autoantibodies across multiple autoimmune disorders^2^. The presence of autoantibodies may precipitate initiation of autoimmune disease prior to the formation of immune complexes and subsequent pathology^15,16^. While autoantibodies are known to be present in individuals with autoimmune disease, they are also found in healthy individuals^17^. However, there is a lack of understanding of the relationship between *HLA* variation and the presence of specific autoantibodies. Here, we examined the extent to which *HLA* genotype influences the autoantibody repertoire in healthy individuals and show that distinct autoreactivity profiles are predictive of HLA class II genotype.

## Methods

### Subjects

Our analysis included serum samples from 741 healthy individuals who volunteered as hematopoietic stem cell donors stored in the NMDP (formerly National Marrow Donor Program) Research Sample Repository (ClinicalTrials.gov protocol NCT04920474). All individuals consented to research and publication of research results and had been previously genotyped for *HLA* class I and class II. These were convenience samples selected based on homozygosity for *HLA-DRB1*, and where possible across the class II region, in order to reduce confounding in the interpretation of the results. Detailed donor demographic information is shown in Supplemental Tables 1 and 2.

The donors included in the study presented a total of 14 *HLA-DRB1* high-resolution (two-field) genotypes. In order to increase statistical power, individuals that shared *HLA* genotype at the first field were merged as a single group (Figure 1). For instance, *HLA-DRB1*15:01* and *HLA-DRB1*15:03* were grouped into a single *HLA-DRB1*15* group. In this manner, five common *HLA-DRB1* allelic groups were chosen as core analysis groups based on sufficient sample size. With a count of 192 subjects, *HLA-DRB1*15* represents the largest core group. The four other groups were *HLA-DRB1*07:01, HLA-DRB1*04* (*HLA-DRB1*04:01, HLA-DRB1*04:04*, and *HLA-DRB1*04:07*), *HLA-DRB1*03:01*, and *HLA-DRB1*01* (*HLA-DRB1*01:01* and *HLA-DRB1*01:02*), with sample sizes of 123, 69, 148, and 96, respectively. The remaining 113 samples were included in the analysis as additional samples to compare against the core groups, which include *HLA-DRB1*11:01, HLA-DRB1*11:04, HLA-DRB1*13:01, HLA-DRB1*13:02*, and *HLA-DRB1*08:02*.

**Figure 1:**
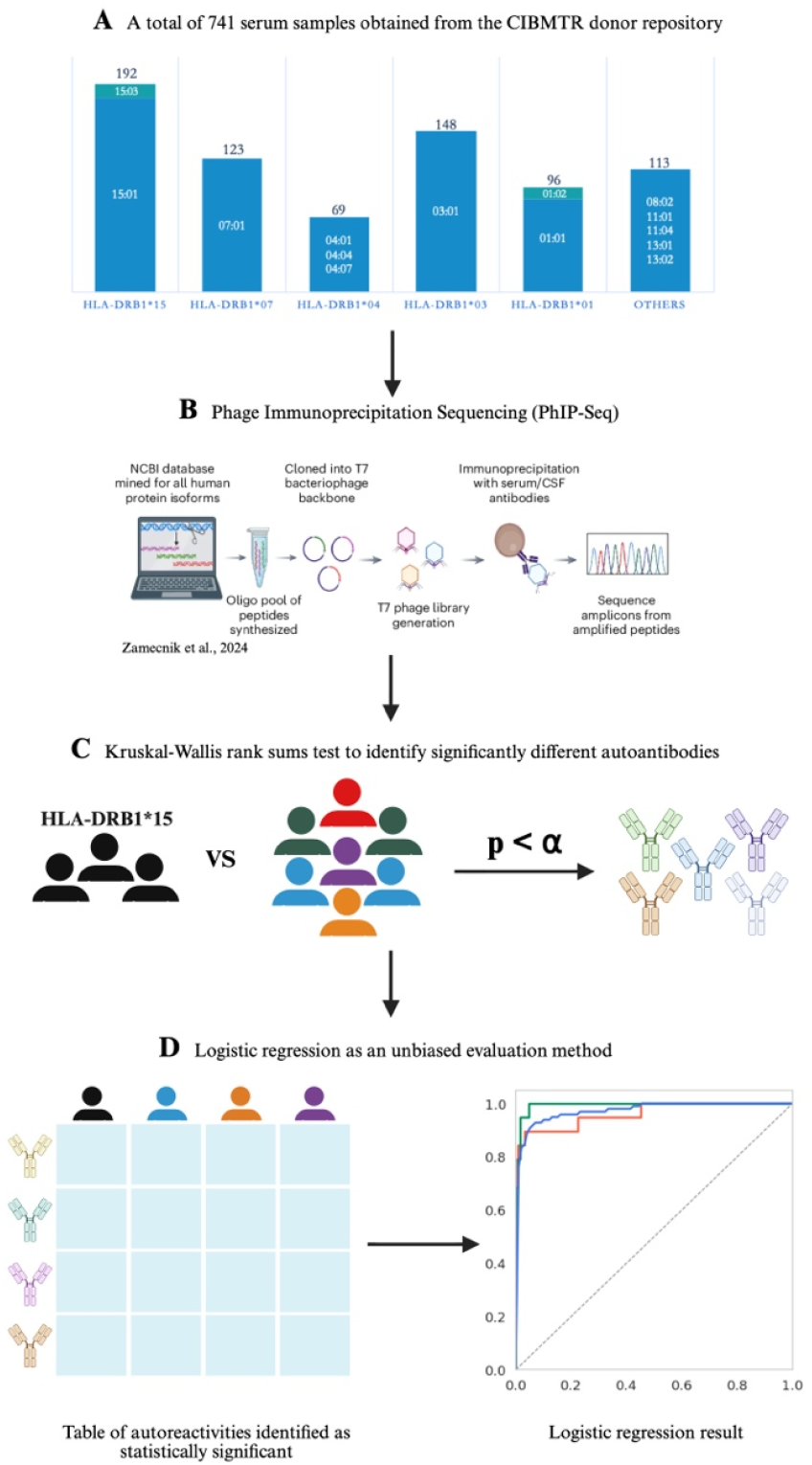
An outline of this study’s design. **A)** serum retrieval of healthy individuals from the CIBMTR donor repository, **B)** Phage Immunoprecipitation Sequencing (PhIP-Seq) performed on the CIBMTR healthy cohort, **C)** Kruskal-Wallis rank sums test to identify autoantibodies that have different reactivity among different genotypes, **D)** logistic regression as means of evaluation.

### Phage Immunoprecipitation Sequencing (PhIP-Seq)

We employed a programmable phage display assay comprised of 731,000 peptides tiled across the human proteome, representing the entire human peptidome, termed PhIP-Seq^18–20^. To generate the human peptidome, we computationally divided the entire human proteome, including all known structural and splice variants, into oligonucleotide sequences encoding 49 amino acid peptides with 24-residue overlap. The phage library was generated using classic molecular cloning such that individual phage particles contain 5-15 copies of a single human peptide expressed on the surface capsid protein^21^.

PhIP-Seq was performed using a high-throughput protocol, which starts with deep-well polypropylene 96-well plates, where the phage library was exposed to 1□µl of human sera in 1:1 storage buffer. The deep-well plates with library and sample were incubated overnight at 4°C to facilitate antibody-phage binding. Beads were added to each reaction well, and a sequence of washes was performed. The immunoprecipitated solution was resuspended and amplified. Filtered solution was transferred to a new pre-blocked deep-well plate where individuals’ sera was added and subjected to another round of immunoprecipitation and amplification, followed by next-generation sequencing (NGS) library prep. Phage DNA from each sample was barcoded and amplified (Phusion PCR, 18 rounds) and subjected to NGS on an Illumina NovaSeq instrument at an average read depth of 1 million reads per sample. Fully detailed PhIP-Seq protocol is described elsewhere^21,22^.

### Binding quantification and peptide linear ranking

Raw sequencing reads from the PhIP-Seq assay were subsequently aligned to a reference library using Bowtie2 to obtain the read count for each sample to each peptide^21,23^. Normalized read counts were then calculated by multiplying the read proportion of each peptide in a sample with 100,000. This operation transformed the raw read count to number of reads per kilo (RPK), meaning that the normalized reads would sum up to 100,000 across the human peptidome. To normalize samples against background, fold change (FC) over mock-IP was calculated by dividing the normalized read by the mean percentage of all bead-only controls in the same peptide. Once the RPK and FC were calculated, we applied a set of filters (MIN_RPK = 0, FC_THRESH1 = 10, FC_THRESH2 = 100, SUM_RPK_THRESH = 50) intended to reduce noise for downstream analysis, yielding us with 102,716 peptides. The data were then transformed into binary hits, using another set of filters to ensure that the gene in which the peptide resides needs to have 1) a summed RPK of 50 or above, and 2) a FC of 100 or above.

### Autoreactivity analysis

To determine which peptide enrichments were statistically significantly different between one genotype group against the others, we used the Kruskal-Wallis rank sums test^24^. To adjust for multiple comparisons, Bonferroni correction was applied such that a p-value threshold of 0.05/6 was enforced. There were a total of six in our analysis: the five primary genotype groups (*HLA-DRB1*15, HLA-DRB1*07, HLA-DRB1*04, HLA-DRB1*03*, and *HLA-DRB1*01*), and one additional group representing all other genotypes.

### HLA genotype prediction from autoreactivity profiles

To test whether *HLA* genotype could be predicted from autoreactivity profiles, we employed multinomial and binary logistic regression. The multinomial model predicts whether a sample belongs to one of five genotype groups: *HLA-DRB1*15, HLA-DRB1*07, HLA-DRB1*04, HLA-DRB1*03*, or *HLA-DRB1*0*1. The binary logistic regression model predicts whether a sample has a specific *HLA-DRB1* genotype. Thus, a total of five binary logistic models were employed: *HLA-DRB1*15* vs rest, *HLA-DRB1*07* vs rest, *HLA-DRB1*04* vs rest, *HLA-DRB1*03* vs rest, and *HLA-DRB1*01* vs rest.

All regression models were performed using only peptides that were significantly different according to the Kruskal-Wallis rank sums test and were exclusive to each genotype group. Due to sample size imbalance between each group against the rest of the samples, we balanced the model during the training process by under-sampling groups that were overrepresented. This was done by using the *class_weight = “balanced”* argument in Scikit-learn’s LogisticRegression function^25^.

For each binary logistic regression, we ran five-fold cross validation to avoid overfitting. Three metrics that we reported to assess the model are accuracy, AUC-ROC (Area Under the Receiver Operating Characteristic Curve), and F1 score. The accuracy can be calculated by dividing the true positive (TP) and true negative (TN) with the total samples 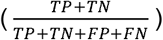. The AUC-ROC was computed by plotting the true positive rate (TPR) against the false positive rate (FPR) across different thresholds and then calculating the area under this curve to assess the overall model discrimination. The F1 score is particularly informative because it reveals if class imbalance is causing the model to disproportionately predict one class. The F1 score for a particular class was calculated using Precision 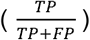 and Recall 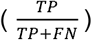 with the following formula: 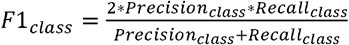, where a class is either *HLA-DRB1*01, HLA-DRB1*03, HLA-DRB1*04, HLA-DRB1*07*, or *HLA-DRB1*15*. All three metrics were calculated using Scikit-learn^25^.

To understand what drives the model’s decision, we ranked the binary logistic regression coefficients to find features (here, specific peptide enrichments) that were the most influential. We ranked the absolute value of the coefficients to better understand the magnitude of the features, while keeping track of its direction. Since the model ran through five-fold cross-validation, we ranked the coefficients based on the average coefficients across five iterations.

### Multidimensionality reduction analysis

We performed the Uniform Manifold Approximation and Projection (UMAP)^26^ on the five main *HLA-DRB1* genotype groups. A total of 9,358 peptides were used in this UMAP, consisting of peptides that passed through the Kruskal-Wallis rank sums test and were exclusive to each genotype group. The binary table was first scaled using the StandardScaler() function so that each feature had zero mean and unit variance. Subsequently, the following arguments were used within Python’s UMAP function: n_neighbors = 15, min_dist = 0.0, n_components = 2, metric = cosine.

## Results

### Autoreactivity profiles vary by HLA-DRB1 genotype

Using the Kruskal-Wallis rank-sum test to compare peptide enrichments between each *HLA-DRB1* group versus all others (*HLA-DRB1*01, HLA-DRB1*03, HLA-DRB1*04, HLA-DRB1*07*, and *HLA-DRB1*15* vs. the rest), we identified 1,029, 1,297, 5,691, 909 and 453 self-peptides, respectively, that were significantly enriched in each group relative to the others. Most autoreactivities identified in specific *HLA-DRB1* groups were private, with very few peptides enriched in two or more *HLA-DRB1* groups (Figure 2A). This demonstrates that the autoreactivity profiles in healthy individuals with different *HLA-DRB1* genotypes are generally distinct.

**Figure 2:**
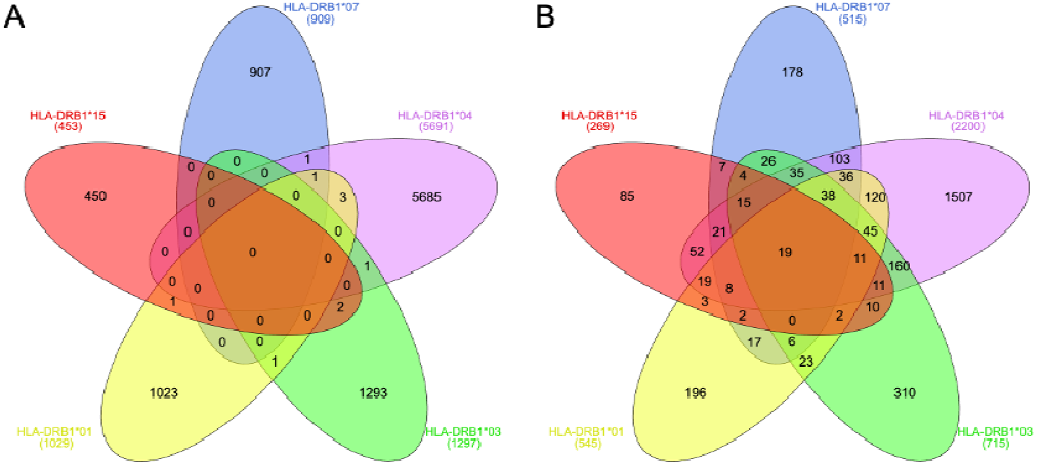
Venn diagrams showing sharing of autoreactivity profiles across genotype groups. **A)** Number of enriched peptides significantly expressed in each *HLA-DRB1* group. **B)** Number of gene/proteins with peptides that were enriched in each *HLA-DRB1* group.

Considering that the peptidome library was built with tiled peptides with 25-mer overlap, it is expected that individuals sharing an *HLA-DRB1* genotype with autoantibodies to a specific peptide may also have antibodies that enrich adjacent, overlapping peptides from the same protein. As an example, *ESPN_526263* and *ESPN_526264*, both originating from the *ESPN* gene, are adjacent to one another and both react to *HLA-DRB1*15* samples. To capture protein-level information from our analysis, we merged all enriched peptides for a given protein in each genotype group. When considering protein-level autoreactive specificity, we found substantially more sharing between genotype groups (Figure 2B). This demonstrates that individuals with different *HLA-DRB1* may have autoantibodies to the same protein (Figure 2B), but enrich distinct peptides (Figure 2A). Nevertheless, we highlight that the majority of protein-level specificity remained exclusive to each *HLA-DRB1* group (196 for *HLA-DRB1*01*, 310 for *HLA-DRB1*03*, 1507 for *HLA-DRB1*04*, 178 for *HLA-DRB1*07* and 269 for *HLA-DRB1*15*). The list of significantly different autoreactivities (peptides and protein) alongside their hit counts can be found in Supplemental Table 3.1 to 3.5.

### Autoreactivity repertoire is predictive of HLA class II genotype

As an unbiased measure of whether the autoreactivity repertoire can be used to predict *HLA* class II genotype, we employed binary and multinomial logistic regression using enriched peptide specificity as predictors. In the binary model, enriched peptide specificities with the following criteria served as predictors: 1) they must be statistically significantly different relative to other groups, and 2) they must be exclusive to the group of interest. As an example, in the *HLA-DRB1*15 vs*. rest model, the matrix contained 1,023 peptides that satisfied these criteria across 628 samples, including *HLA-DRB1*15* (192), *HLA-DRB1*07* (123), *HLA-DRB1*04* (69), *HLA-DRB1*03* (148), and *HLA-DRB1*01* (96). While the model’s accuracy and AUC-ROC score are informative metrics to report, we also present the F1 score to ensure that our models are not biased to one class due to data imbalance.

For the binary model, all groups presented an AUC ranging from 0.96 to 0.98 after undergoing 5-fold cross validation (Figure 3). The *HLA-DRB1*01* group specificities performed best in predicting whether a sample has an *HLA-DRB1*01* genotype (F1_*HLA-DRB1*01*_ = 0.85) from the autoreactivity repertoire. In contrast, *HLA-DRB1*04* specificities performed poorly in predicting whether a sample had an *HLA-DRB1*04* genotype (F1_*HLA-DRB1*04*_ = 0.13). This effect was more apparent from the confusion matrix, shown in Supplemental Figure 1, where the model only correctly predicted one out of fourteen *HLA-DRB1*04* samples, albeit with satisfactory AUC and accuracy. We hypothesized that the poor predictive performance was likely due to the small sample size of the HLA-DRB1*04 group (n = 69), which may have caused the Kruskal–Wallis rank-sum test to identify a disproportionately large number of significantly different enriched peptides (5,691 in this case—several orders of magnitude higher than the average of 922 in other groups).

**Figure 3:**
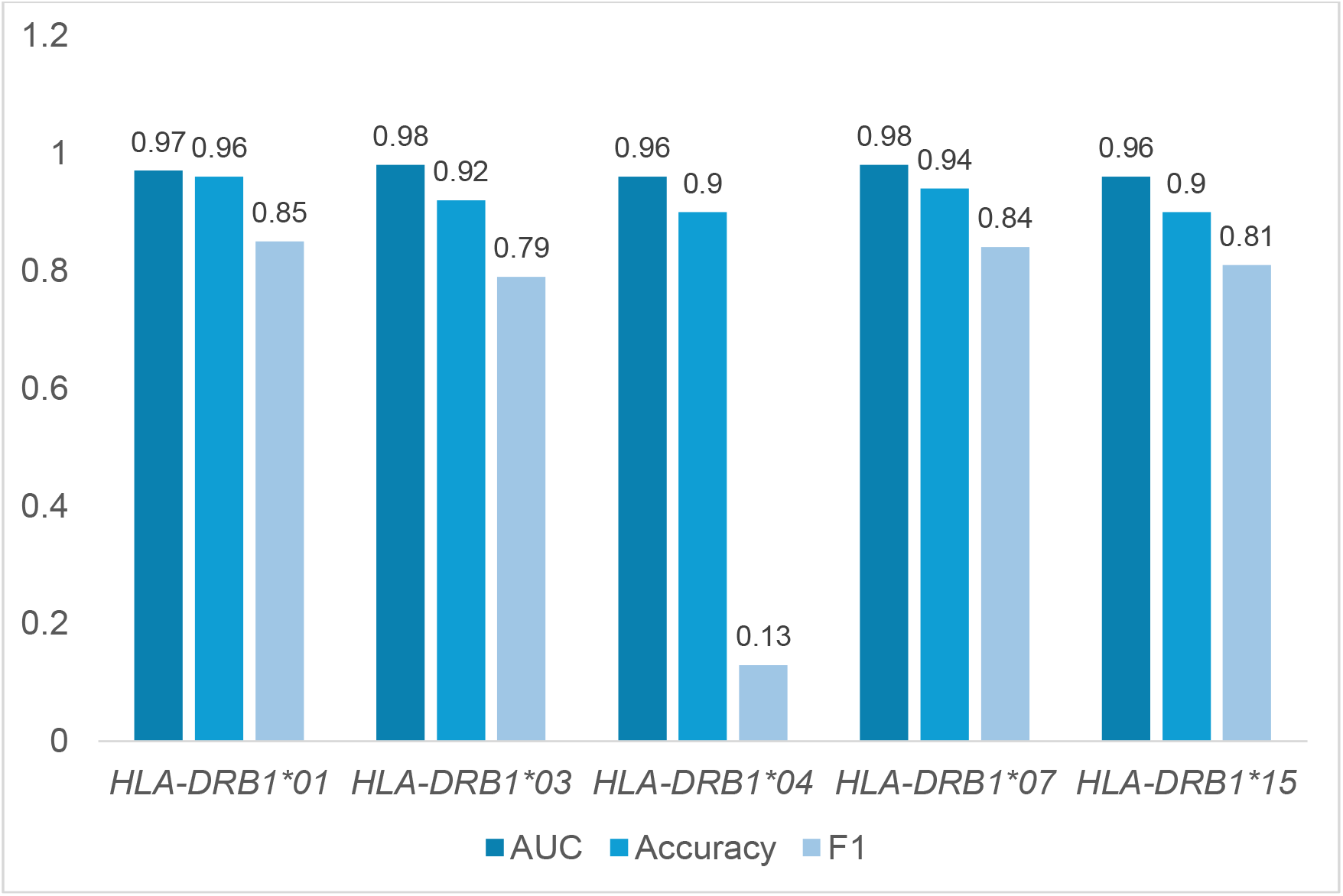
Binary logistic regression performance metrics, which includes the AUC-ROC, accuracy, and F1 score. The models successfully determined a sample’s genotype according to autoreactivity, as shown by an average AUC of 0.97 and an average accuracy of 0.92 across the groups. The F1 score for *HLA-DRB1*04* is particularly low due to its sample size being smaller than the rest of the groups. This caused the model to preferentially predict the majority group (non-*HLA-DRB1*04*), even after balancing the training data.

To understand whether low sample count for the *HLA-DRB1*04* group was underlying poor performance in that group, we conducted a sensitivity analysis based on sample size, and randomly removed samples from *HLA-DRB1*15* to match the number *HLA-DRB1*04* samples. Running Kruskal-Wallis rank sums test between 69 *HLA-DRB1*15* samples and the 549 remaining yielded 6,372 enriched peptides that were statistically significantly different, similar in magnitude to *HLA-DRB1*04*. We also performed binary logistic regression with these 618 samples and 6,372 peptides, which obtained an accuracy and AUC of 0.89 and 0.99. However, much like *HLA-DRB1*04*, we observed an F1_*HLA-DRB1*15*_ value of 0.13, indicating that the model did not perform optimally due to the class imbalance.

To understand which features were driving decisions in the binary model, we ranked the coefficients that represent the importance of each enriched peptide. These coefficients can either be a positive predictor (shown in blue, Figure 4), or a negative predictor (shown in red, Figure 4). A positive predictor is a peptide that shifts the model’s decision towards categorizing a sample to the genotype of interest, whereas a negative predictor pushes that decision in the opposite direction. For example, in the *HLA-DRB1*03* logistic regression model, enrichment of peptide *NHSL2_57958* functions as a positive predictor (blue box), while *MAML3_311538* serves as a negative predictor (red box). Thus, enrichment of *NHSL2_57958* increases the likelihood that a sample will be classified as *HLA-DRB1*03*, whereas enrichment of *MAML3_311538* pushes the classification toward non-*HLA-DRB1*03*. The complete ranking of predictors for our model is given in Supplemental Table 4.1 to 4.5.

**Figure 4:**
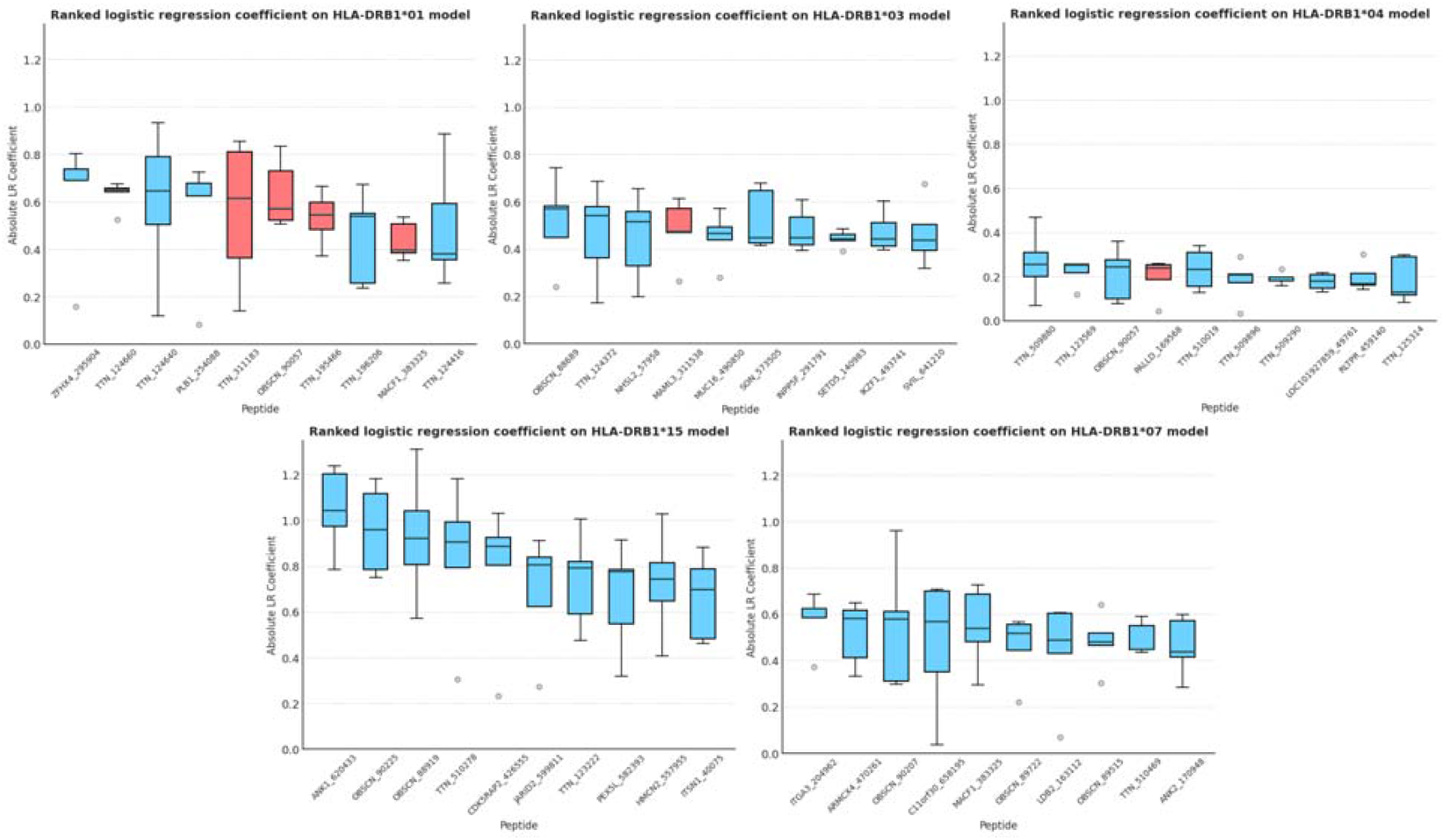
Ranked binary logistic regression coefficients, based on five iterations of cross-validation. Each box in the boxplot represents the feature’s coefficient from five iterations of five-fold cross-validation. The colors of the boxes indicate whether an enriched peptide acts as a positive or negative predictor. In this context, a positive predictor is a feature (an enriched peptide) that shifts the model’s decision toward a positive outcome.

Multinomial logistic regression returned an accuracy of 76% when the five *HLA-DRB1* groups were tested (Figure 5A). The *HLA-DRB1*04* group showed an F1 score of 0.55 while the F1 score ranged from 0.75 to 0.81 for the remaining four groups. When we performed a multinomial logistic regression excluding *HLA-DRB1*04* group, we observed an accuracy improvement to 90% (Figure 5B). The F1 score ranged from 0.89 (*HLA-DRB1*01* and *HLA-DRB1*03*) to 0.93 (*HLA-DRB1*07*). These results support the notion that a given *HLA-DRB1* genotype can be predicted from a given autoreactivity repertoire, owing to distinct specificity profiles.

**Figure 5:**
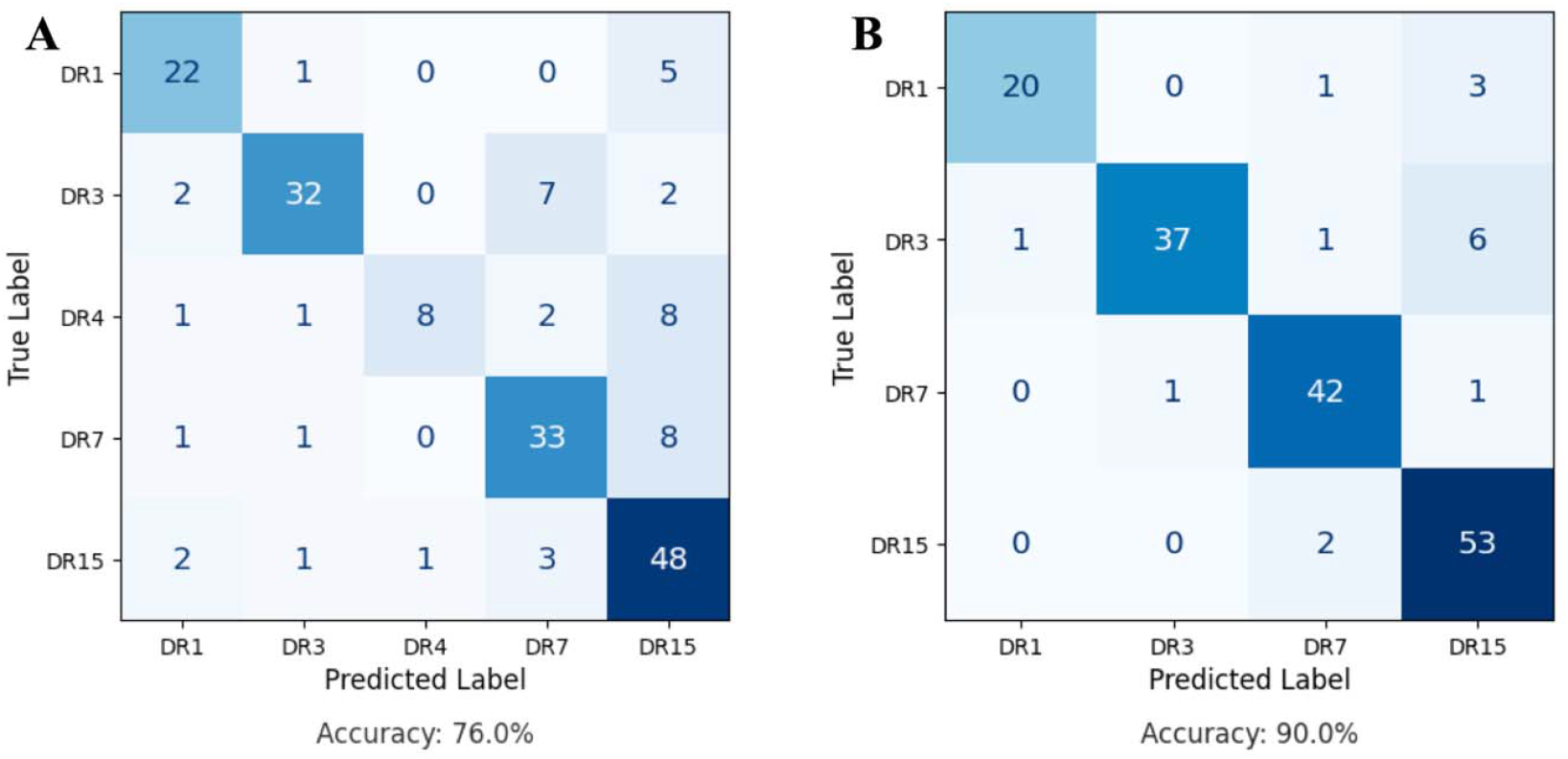
Confusion matrix from the resulting multinomial logistic regression. **A)** Multinomial logistic regression including all five *HLA-DRB1* groups. **B)** Multinomial logistic regression excluding *HLA-DRB1*04* group.

### Autoreactivity profiles cluster donors according to HLA-DRB1 genotype

To examine whether *HLA* class II genotype groups cluster according to autoreactivity profiles, we applied UMAP to reduce the dimensionality of our data (628 samples × 9,358 peptides) into a 2D representation, as shown in Figure 6. By color-coding each point according to the sample’s *HLA-DRB1* genotype, we observed a clear tendency for samples with the same genotype to cluster together, indicating genotype-specific patterns in autoreactivity. Unsurprisingly, *HLA-DRB1*04* exhibited weaker clustering performance compared to the remaining groups, which was likely a consequence of its small sample size. Nevertheless, our results clearly demonstrate that HLA class II genotypes cluster according to autoreactivity profiles.

**Figure 6:**
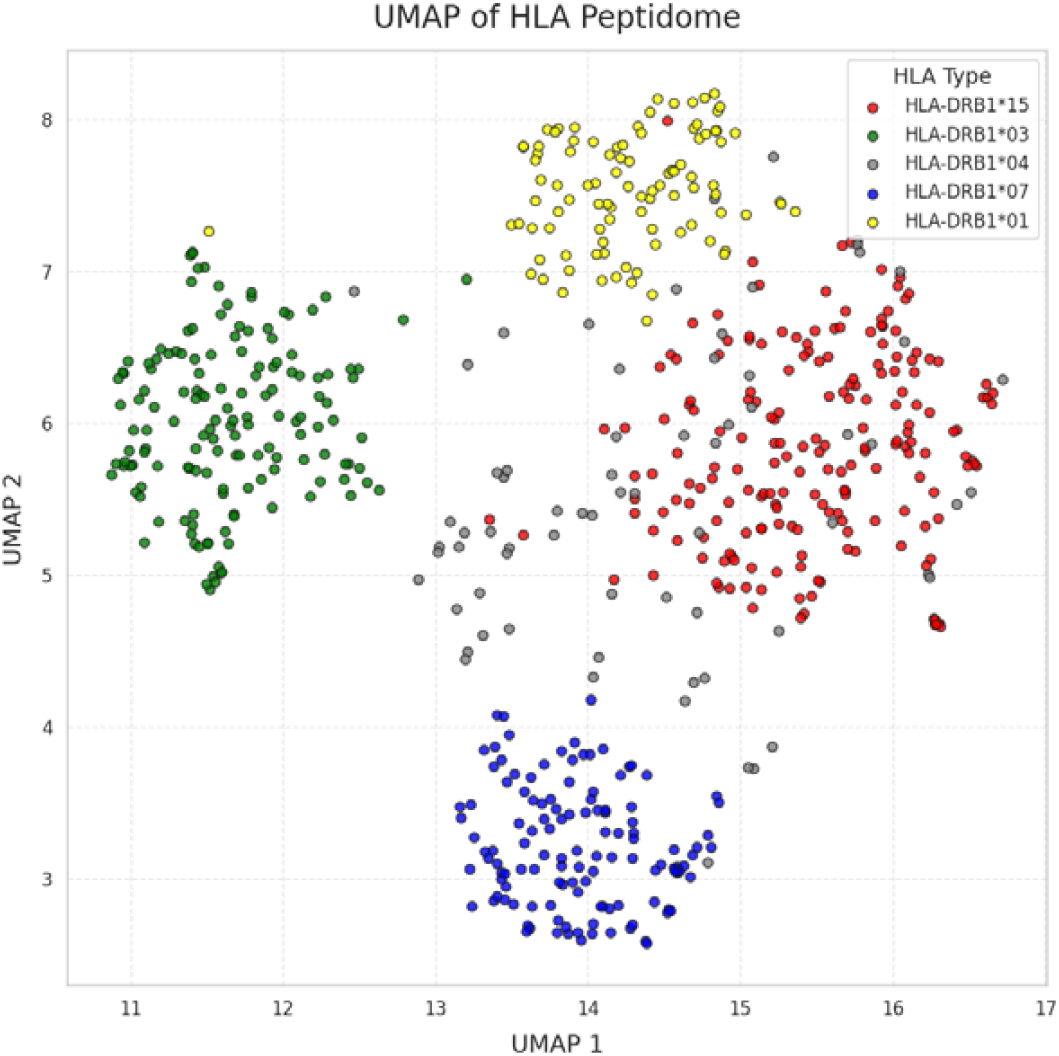
UMAP of autoantigen enrichments of *HLA-DRB*1 groups. Each point in this plot is a sample with a known genotype. A total of 9,358 peptides were represented in this UMAP, consisting of enriched peptides that passed through the Kruskal-Wallis rank sums test and were exclusive to each genotype group.

## Discussion

Previous studies have demonstrated the presence of autoantibodies in disease-free individuals^16,17,21,34,35^. Further, it has been shown that individual IgG autoantibody profiles in the blood are stable over long periods of time^17,36,37^, there is no gender bias^35^, and healthy individuals share common autoantibodies^34,35^. In a study with twins, Venkataraman et al (2022) demonstrated MHC class II molecules strongly influence the binding specificities of antibodies targeting EBV (Epstein-Barr virus) and its antibody epitope selection as a heritable trait^14^. Haller-Kikkatalo *et al*., (2017) conducted autoantibody profiling in disease-free individuals and concluded that presence of tissue non-specific autoantibodies might serve as prognostic markers for future disease. Given the role of autoantibodies and the known association of *HLA* class II variation with autoimmune disease risk, in the present study we sought to understand whether *HLA* class II genotype influences the autoreactome in the absence of overt autoimmune disease. We examined 741 serum samples from healthy *HLA-DRB1* homozygous donors for autoreactive specificity against the entire human proteome, consisting of over 731,000 peptides, demonstrating that autoreactivity profiles are differentially enriched between the five *HLA-DRB1* groups analyzed.

The PhIP-Seq platform described here has previously been used to identify specific autoantibody specificities in a disease setting^38^. For instance, in a case-control study, Zamecnik *et al*., (2024) found an autoantibody signature that is predictive to multiple sclerosis (MS)^21^, while Bodansky *et al*., (2024) carried out a similar comparison with multisystem inflammatory syndrome in children (MIS-C)^39^. Unlike these case-control comparisons, our model emphasizes variation within the healthy population to show the influence of the *HLA* class II towards the autoreactive profile in the absence of diagnosed autoimmune disease.

Despite our finding that *HLA* class II shapes the autoreactome, the mechanism of this process remains unclear. *HLA* class II alleles determine which peptides are presented to T cells, leading to their activation and the provision of T cell help. This process triggers cytokine release, which stimulates B cells to produce antibodies—or, in cases of self-peptide presentation, autoantibodies. To prevent self-peptide presentation, autoreactive T cells are typically eliminated via clonal deletion in the thymus during its development^40^. This process, however, is imperfect and autoreactive T cells often escape clonal deletion^40^.

To the best of our knowledge, this is the first study to showcase that the autoreactome in healthy individuals is shaped by *HLA* class II genotype. Further, through a supervised machine learning regression model we were able to demonstrate that the autoreactivity profiles can be used to predict *HLA-DRB1* genotype, reaching a performance accuracy of up to 90% for a multinomial model and 96% for a binomial model (*HLA-DRB1*01*). This work has important implications for understanding both the role of HLA in disease predisposition, and the immunological conditions that precede overt autoimmune disease. Future studies in more diverse cohorts and including other *HLA* genotypes will provide additional evidence and clarity regarding the role of *HLA* variation in shaping the autoreactome in healthy people.

