## Supplemental Figure 1 for "Autoreactomes in Healthy Individuals Vary According to HLA Class II Genotype"

**Supplemental Figure 1** – ROC curve and confusion matrix for five binary logistic regression models for *HLA-DRB1*01, HLA-DRB1*03, HLA-DRB1*04, HLA-DRB1*07,* and *HLA-DRB1*15*

| 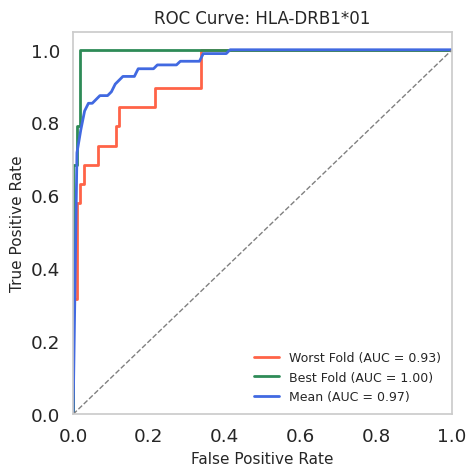 | 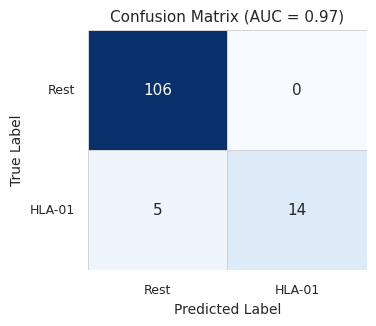 |
| --- | --- |
| 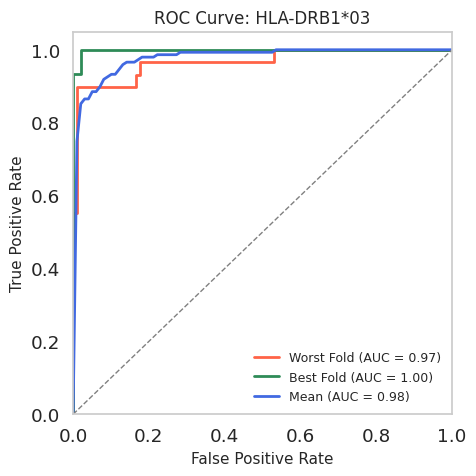 | 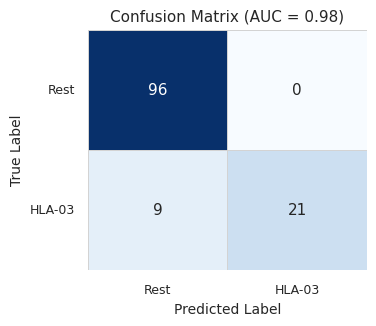 |
| 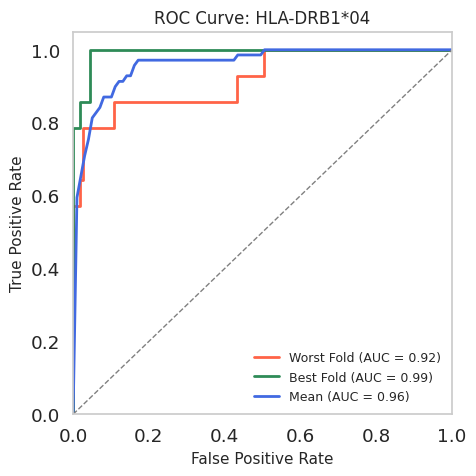 | 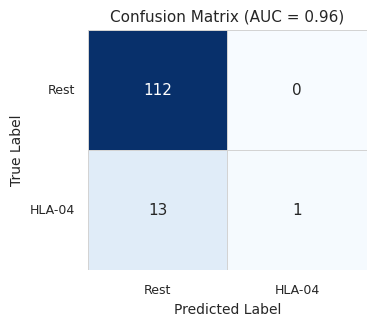 |
| 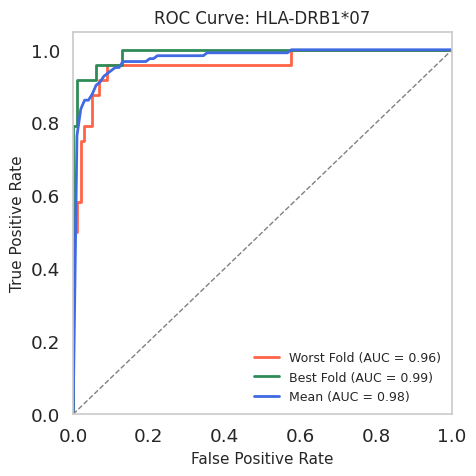 | 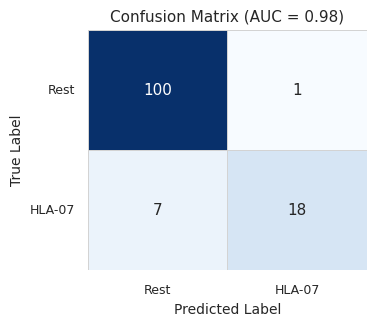 |
| 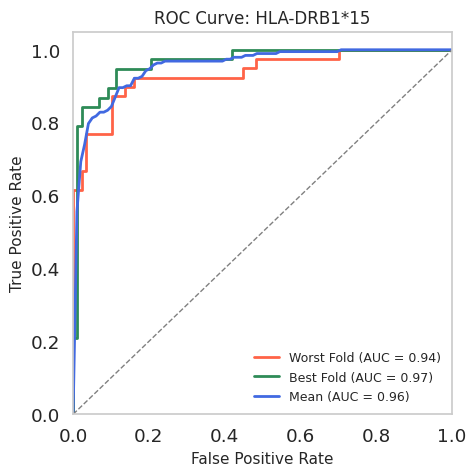 | 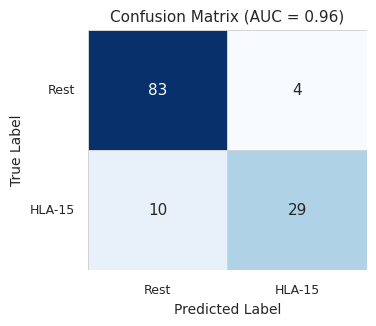 |
