## Supplemental Table 1 for "Autoreactomes in Healthy Individuals Vary According to HLA Class II Genotype"

| **Supplemental Table 1**. Sex and age distribution | | | | | | |
| --- | --- | --- | --- | --- | --- | --- |
| ***HLA* group** | **Female N (%)** | **Male N (%)** | **Total N** | **p-value*** | **Mean age (SD)** | **p-value**** |
| *DRB1*01* | 35 (36.5) | 61 (63.5) | 96 | 0.227 | 35.3 (9.8) | 0.553 |
| *DRB1*03* | 37 (24.8) | 112 (75.2) | 149 | 0.014 | 33.3 (9.6) | 0.058 |
| *DRB1*04* | 27 (38.6) | 43 (61.4) | 70 | 0.163 | 34.8 (9.1) | 0.914 |
| *DRB1*07* | 34 (27.2) | 91 (78.8) | 125 | 0.094 | 34.9 (10.2) | 0.832 |
| *DRB1*15* | 62 (32.3) | 130 (57.5) | 192 | 0.493 | 33.4 (9.8) | 0.032 |
| Other *DRB1* | 48 (32.6) | 65 (67.4) | 113 | 0.017 | 38 (9.6) | <0.001 |
| Total | 243 (32.6) | 502 (67.4) | 745 | - | 34.7 (9.8) | - |
| *Fisher exact test, **t-test | | | | | | |
