## Supplemental Table 2 for "Autoreactomes in Healthy Individuals Vary According to HLA Class II Genotype"

| **Supplemental Table 2**. Ancestry/Race distribution | | | | | | | | | | | | | | |
| --- | --- | --- | --- | --- | --- | --- | --- | --- | --- | --- | --- | --- | --- | --- |
| ***HLA* group** | **Ancestry/Race** | | | | | | | | | | | | | **Total** |
|  | **AFA** | **API** | **CARB** | **CARIBI** | **EUR** | **DEC** | **HIS** | **MSWHIS** | **MULTI** | **NAM** | **UNK** | **WCARIB** | **WSCA** |  |
| *DRB1*01* | 0 | 0 | 0 | 0 | 81 | 1 | 1 | 0 | 1 | 0 | 11 | 0 | 1 | 96 |
| *DRB1*03* | 2 | 5 | 0 | 0 | 123 | 0 | 2 | 0 | 4 | 0 | 13 | 0 | 0 | 149 |
| *DRB1*04* | 0 | 0 | 0 | 0 | 60 | 0 | 3 | 0 | 2 | 1 | 3 | 1 | 0 | 70 |
| *DRB1*07* | 4 | 4 | 0 | 1 | 92 | 0 | 4 | 0 | 3 | 5 | 11 | 0 | 1 | 125 |
| *DRB1*15* | 13 | 2 | 1 | 0 | 150 | 0 | 1 | 2 | 5 | 0 | 18 | 0 | 0 | 192 |
| Other *DRB1* | 0 | 1 | 0 | 0 | 82 | 0 | 13 | 1 | 1 | 1 | 14 | 0 | 0 | 113 |
| Total | 19 | 12 | 1 | 1 | 588 | 1 | 24 | 3 | 16 | 7 | 70 | 1 | 2 | 745 |
| AFA: African American; API: Asian/Pacific Islander; CARB: Black Caribbean; CARIBI: Caribbean Indian; EUR: European descent; HIS: Hispanic; MSWHIS: Mexican or Chicano; MULTI: multiple ancestry; NAM: Native American; UNK: unknown ancestry; WCARIB: White Caribbean; WSCA: White South or Central America. | | | | | | | | | | | | | | |
